# Assessment of drive efficiency and resistance allele formation of a homing gene drive in the mosquito *Aedes aegypti*

**DOI:** 10.1101/2024.09.24.614707

**Authors:** Xiaozhen Yang, Xuejiao Xu, Yixian Chen, Jiajia Wei, Wanting Huang, Songqing Wu, Jackson Champer, Junxiang Wang

## Abstract

*Aedes aegypti*, known for transmitting viruses such as dengue, zika, and yellow fever, poses a significant public health threat. Conventional insecticides give rise to a range of issues, including ecological contamination and insect resistance. Hence, there is a pressing demand for environmentally friendly, safer, and more efficacious strategies for mosquito control. With the rapid advancement of the CRISPR/Cas9 system in gene function exploration and pest population control, substantial progress has been achieved in utilizing CRISPR/Cas9-based gene drive systems across various mosquito species. Only a few studies on gene drive technology have been conducted in *A. aegypti*. In this study, we constructed two complete drives for *A. aegypti* with different Cas9 promoters, each targeting *kmo*. Our drive based on *Pub*-Cas9 had limited activity, but one with *exu*-Cas9 exhibited super-Mendelian inheritance rates of approximately 60%. We observed low but detectable somatic activity of the drive and no evidence of maternally deposited Cas9. Germline resistance allele formation rates were similar to drive conversion rates, but most wild-type alleles in the germline remained uncut. Injections into the *exu*-Cas9 drive line had 100% knockout efficiency among surviving offspring at three separate target genes. These results support the development and application of novel genetic pest control technologies aimed at combating *A. aegypti*.

## Introduction

*Aedes aegypti* is the primary vector of several highly pathogenic viruses including dengue, Zika and yellow fever, which pose a major threat to human health^1, 2^. Currently, there is no effective vaccine for preventing these infectious diseases. Thus, effective control of the *Aedes* vector mosquito population remains the principal approach for prevention. Conventional, pesticide-based strategies have been shown to lead to resistance and subsequent decline in efficiency^3^, and there is a race between the development of new pesticides and of pesticide resistance. As pesticide usage increases, resistance development pressure also increases, leading to a rapid decline in effectiveness. Consequently, it is imperative to explore alternative pest control strategies that are safe, efficacious, and environmentally sustainable. Recently, many studies have introduced transgenic elements into insects’ genome with the aim of achieving “insect treatment through insects” using mechanisms such as gene drive^4, 5^.

Gene drive is the phenomenon of super-Mendelian inheritance (>50%) of an allele in the next generation^4^. This mechanism can be applied for population suppression and population modification. Among multiple possible gene drive approaches, CRISPR/ Cas9-based homing drive is the most well-studied and perhaps the most powerful^6, 7^. In this system, Cas9 and guide RNA (gRNA) are inserted into the genome, and the expressed gRNA guides Cas9 to cleave a wild-type allele of the target gene, resulting in a DNA double strand break^8^. During DNA repair, cells utilize the drive allele as a template for homology-directed repair, achieving the transformation from heterozygous to homozygous for the drive, thereby increasing the chance of inheritance among the offspring. This process is called “homing” or “drive conversion.” However, when end-joining repair is applied in lieu of homology-directed repair, mutations involving indels can form, leading to the failure of gRNA to recognize the cutting site. These mutations are called a resistance allele^9^. The accumulation of resistance alleles can reduce the proportion of drive alleles in a population and even result in gene drive failure.

Homing drives can be divided into two types based on the localization of the endogenous Cas9 and gRNA: standard “complete” drive (Cas9 and gRNA are both inserted into the gRNA site) ^10, 11^ and split drive (one of these, usually Cas9, is not inserted into the driving site)^12, 13^. To date, homing drive systems of both types have been established in various insect species, including *Drosophila melanogaster*^14–29^, multiple mosquito species (*Anopheles gambiae*^11, 30, 31^, *Anopheles stephensi*^10, 32, 33^, *Anopheles Coluzzii*^31^*, Aedes aegypti*^34–38^, *Culex quinquefasciatus*^39, 40^), and *Plutella xylostella*^41^. In both *Anopheles* species, the transmission rates of drive allele were often close to 100%^10, 11, 30–32^. A modification gene drive in *An. stephensi* exhibited a transfer efficiency of ≥95% within 5-11 generations during cage trials^32^. For *An. gambiae*, a modification gene drive carrying anti-plasmodium cargo genes demonstrated a high drive conversion efficiency (≥98%) while reducing malaria incidence by ≥90% within 3 months according modeling^31^. Another suppression drive also reached ≥95% transmission rate targeting *doublesex* gene in *An. gambiae* ^30^. In *Ae. aegypti* a series of drives have been established to assess the drive efficiency of Cas9 expression through various endogenous promoters. Several exhibited inheritance rates ranging from 50% to 70%^34–38^. Anderson *et al*. discovered that among the *sds3* promoter lines cross to a 4-gRNA element targeting *kmo*, one line, together with some Cas9 lines based on the shu promoter, demonstrated high inheritance rates of over 80%, while two other two *sdr3* lines displayed an inheritance rate close to 50%^35^. The study of Verkuijl *et al*. also reported a lower inheritance rate of only 67% for *sds3*-Cas9^36^. Of these, only Reid *et al*. characterized resistance allele formation, which was moderate for the *nanos* promoter and low for the *zpg* promoter in comparison to the drive conversion rate^38^.

In this study, we generated two standard drives in *Ae. aegypti* targeting the *kmo* gene and employing distinct endogenous promoters to drive the expression of Cas9 (PubCas9-*kmo* and ExuCas9-*kmo*^42^). By crossing transgenic strains with wild-type, we compared the inheritance rate of the two transgenic strains. We also assessed resistance allele formation with the *exu* promoter for Cas9. Overall, we found that Pub was not an effective Cas9 promoter and that *exu*-Cas9 produced a moderate number of resistance allele in both males and females.

## Methods

### Mosquito rearing and maintenance

*Ae. aegypti* Haikou wild strain (wild-type strain) was obtained from the Fujian International Travel Health Care Center (Fujian, Fuzhou, China) and reared in laboratory environment for more than 65 generations. All mosquitoes were maintained at 26 ± 1 °C and 83% ± 3% relative humidity with 14h/10h day/night light cycles. Larvae were reared in purified water and fed by goldfish feed (Tetra, Germany). Adults were fed with 10% sucrose solution. Three days after mating, females were fed with sterile defibrillated bovine blood (Maojie, China), and eggs were collected several days later for hatching.

### Plasmids design and assembly

The Cas9 coding sequence was optimized for *Aedes aegypti* codon usage, with a SV40 nuclear localization signal in the N-terminus and a nucleoplasmin nuclear localization signal in the C-terminus. The genome DNA of *Aedes aegypti* was extracted from five 4th instar larvae with the Insect DNA Isolation Kit (OMEGA, USA). Total RNA of *Aedes aegypti* was extracted from five 4th instar larvae by TRIZOL. cDNA was prepared using ReverTra Ace qPCR RT Master Mix with gDNA Remover kit (TOYOBO, Japan). *AeExu*, *AePub*, *AeU6* promoter, *AeU6* 3′ UTR and *kmo* gene homologous arm sequences were amplified from gDNA by Q5 Hot Start High-Fidelity 2X Master Mix (New England Biolabs, UAS) and cloned on Cas9-DsRed backbone.

Plasmids were constructed by In-Fusion Cloning using IN-Fusion HD Cloning Kit (TaKaRa, Japan). Correct clones were confirmed by Sanger sequencing and double enzyme digestion. EndoFree Plasmid DNA was extracted with the EndoFree Plasmid Giga Kit (Qiagen, Germany). Plasmid and target gene sequences are available on GitHub (https://github.com/jchamper/Aedes-Complete-Drive).

### gRNA Expression

To select a U6 promoter for gRNA expression, we conducted a BLAST alignment between sequences from *D. melanogaster* U6 gene (NR002083) and *An. Stephensi* U6 gene (ASTE015697) against the *Ae. aegypti* genome^43^. We identified seven highly conserved U6 genes, among which AeU6-1 (AEEL017774) was known to effectively drive shRNA to induce target gene knockdown in the *Ae. aegypti* cell line ATC-10^43^. However, when using microinjection of U6-1-gRNA and Pub-Cas9 plasmids into *Ae. aegypti* embryos, no gene editing activity was detected^44^. This observation could be attributed to the limited sample size (500-700 injected embryos) utilized in this experiment^44^. It may also suggest that the transcriptional activity of CRISPR/Cas9 mediated by exogenous plasmids is not sufficiently robust in *Ae. aegypti*^44^. Given that the *Pub* promoter has been shown to effectively drive high expression of exogenous genes and TALEN gene editing activity in *Ae. aegypti* embryos, it is possible that the 96 bp AeU61 promoter was too short, lacking crucial regulatory elements^45^. The effective length of U6 promoters driving gRNA expression in *D. melanogaster*, *An. gambiae*, and *An. Stephensi* ranges from 300 to 500 bp^6, 10, 11^. As a result, for the selection of U6 promoter, we opted for the first 400 bases before the start of U6-1 as the promoter driving *kmo*-gRNA expression, including a 300 bp termination sequence derived from U6-1.

### gRNA and dsRNA synthesis

A CRISPR/Cas9 target site in the *kmo* gene was selected, and an RNAi target region of *ku70* gene was also selected to increase injection efficiency. The template dsDNA for the transcription of *kmo*-sgRNA and *ku70*-dsRNA was synthesized using Q5 Hot Start High-Fidelity 2x Master Mix. The template dsDNA was transcribed *in vitro* by MEGAscript T7 Transcription Kit (Ambion, USA), and sgRNA and dsRNA were purified by phenol:chloroform extraction and ethanol precipitation. All gRNAs sequences are shown in Table S1.

### Embryo microinjection

The collection and microinjection of mosquito embryos was performed as described previously^46^. Injection solution contained 300 ng/μL spyCas9 protein (NEB, USA), 100 ng/μL *kmo*-sgRNA, 300 ng/μL ku70-dsRNA and 500 ng/μL PubCas9-*kmo* or exuCas9-*kmo* plasmid. Embryos were injected using Nanojuct Ⅲ^TM^ (Drummond USA). The needle was pulled by vertical needle puller PC-10 (Narishige, Japan) with supporting capillary glass tube and was opened using the needle grinder EG-400 (Narishige, Japan). The injected mosquito eggs were transferred to a wet filter paper and affixed to the wall of the paper cup to remove oil. Cotton saturated with water was placed in the paper cup to maintain humidity. Five days later, the eggs were placed upside down in filtered water to hatch.

### Fluorescence screening

Each G0 male mosquito was mated with five wild-type female mosquitoes and each G0 female mosquito was mated with three wild-type male mosquitoes. After three days, male crosses and female crosses were pooled together (injected males and injected females were maintained in separate batches from each other) to bloodfeed and lay eggs. Three rounds of feeding and egg collection were performed for each batch. The G1-generation mosquitoes were raised until the 4th instar of larva or the pupal stage, and fluorescence screening was performed by stereo-fluorescence microscopy (Leica, Germany). All positive mutants with red fluorescence were collected to establish the line.

### Identification of genotype in *Ae. aegypti*

The genomic DNA of different phenotypes were extracted using 50 μL of chelex-100 buffer with a hind leg of each adult as described previously^47^. According to the sequences on the right and left homologous arm of the *kmo* target site and donor plasmid elements, the genotypes were verified by amplification with specific primers using Primix TaqTM (TaKaRa, Japan).

To detect the cleavage at the *kmo* gRNA site, genomic DNA was extracted from 10 *kmo*- strain larvae. *kmo* gene (about 600 bp with the *kmo* gRNA site) was amplified with Q5 Hot Start High-Fidelity 2X Master Mix (NEB, USA) from *kmo*- genomic DNA. Amplicons were purified using Gel Extraction Kit (OMEGA, USA) and cloned using the CloneJET PCR Cloning Kit (Ambion, USA). Ten clones were then Sanger sequenced.

### Crosses and assessment

A single female or male adult of PubCas9-*kmo* or exuCas9-*kmo* strain was randomly selected to cross with three male wild-type or five female wild-type mosquitoes, respectively. Fluorescence and white eye phenotype were observed at the 4th instar larvae or pupae stages of G1 by fluorescent microscopy, and the number and proportion of each phenotype in G1 were calculated. Female and male G1 with white eyes and normal eyes with red fluorescence were randomly selected to cross with wild-type, and the phenotype of G2 was observed and statistically analyzed. These steps were repeated to determine the phenotypes of G3 and G4.

### Data analysis

Statistical analysis was performed by Excel and GraphPad prism9. Alternate analysis used linear regression with binomial model to account for batch effects as described previously^48^. Drive inheritance was compared to the Mendelian expectation using the Binomial Test. Two groups were compared using Fisher’s exact test.

## Results

### Generation of drive mosquitoes

Our drive constructs consist of four main elements: (1) a Cas9 gene flanked by an endogenous promoter and 5′ UTR of *Aedes aegypti* and an SV40 terminator; (2) a reporter gene, DsRed under the control of the OpIE2 promoter; (3) a gRNA cassette to produce a single gRNA targeting *kmo* that is expressed by the *Ae. aegypti* polymerase III promoter U6-1; (4) two ∼1.5kb homology arms in the donor plasmid matching the flanking genomic sequences of the *kmo* gRNA cut site to promote homology-directed repair-based incorporation of the drive into the genome. To promote high expression of Cas9, we chose two promoters. One is 1392 bp of AAEL003877 (*polyubiquitin*), which has high expression level during many developmental life stages. The other is AAEL010097 (*Exuperentia*), which has high expression level in the ovary after blood feeding. We built two donor plasmids named PubCas9-*kmo* (Fig 1a) and ExuCas9-*kmo* (Fig 1b).

**Fig. 1.**
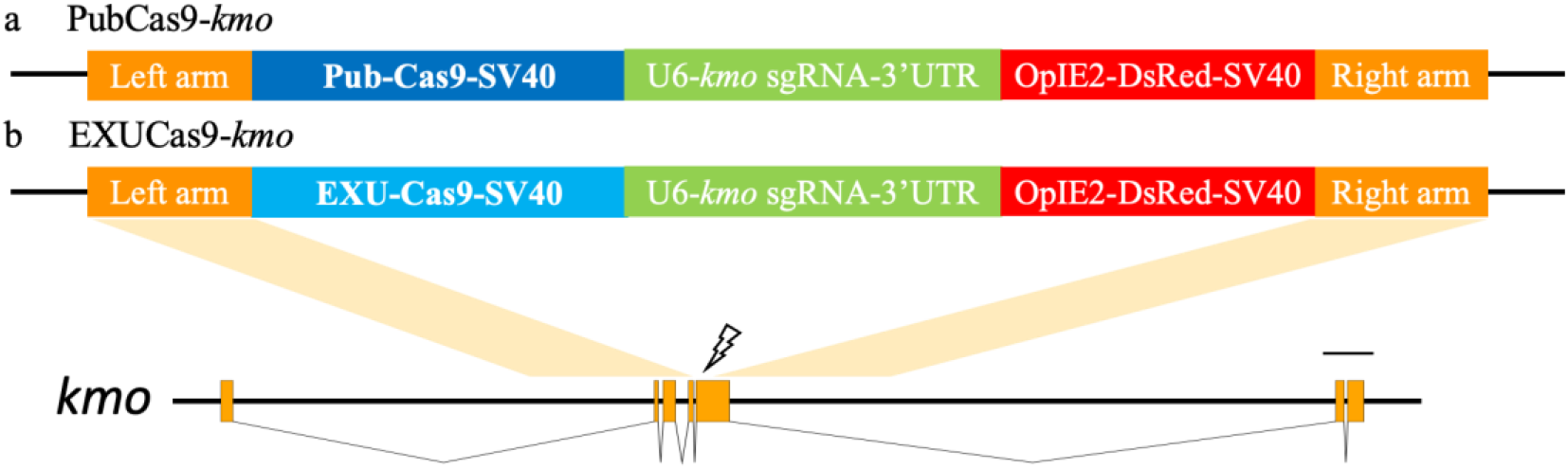
Schematic diagram of the knock-in vector. (**a**) Our donor cassette using the *Pub* promoter to express Cas9. (**b**) Our donor cassette using the *Exu* promoter to express Cas9. (a) and (b) used the same gRNA targeting the exon5 of *kmo* gene. Both donor plasmids contain a DsRed marker.

To make sure that our selected *kmo* gRNA was functional, we performed knockout experiments. The mix of gRNA and Cas9 mRNA was injected into 350 mosquito eggs. A total of 42 larvae (12%) hatched. Finally, 39 larvae developed to adults (Table S2). By observing the eye phenotype, it was found that three mosquitoes (two males and one female) had completely lost their eye pigment, showing a white eye phenotype (Table S2 and Fig S1).

To generate transgenic mosquitoes, Cas9 protein, *kmo*-gRNA and donor plasmid were injected into fresh eggs of a wild-type strain. In addition, dsRNA targeting *Ku70* was added to the injection to reduce the activity of the end-joining repair pathway and thus increase the probability of homology-directed repair. For PubCas9-*kmo*, a total of 435 eggs were injected, resulting in 27 surviving adults (15 males and 12 females) (Table 1). Adults were crossed with wild-type mosquitoes and were assigned to two male-founder and one female-founder pools for blood feeding. After screening 8071 G1 larvae, 254 positive larvae (3.14%) displaying red fluorescence were identified (Table 1). Similarly, for the ExuCas9-*kmo* plasmid, 300 eggs were injected with 21 surviving individuals (13 males and 8 females), ultimately obtaining 283 transgenic G1 individuals out of 5934 total larvae (4.76%) (Table 1).

**Table 1.** CRISPR Injection of PubCas9-*kmo* and ExuCas9-*kmo* Lines.

| Donor | G0<br>Injected<br>Embryos | G0<br>Survivor<br>Batches | G1 Phenotype (Red fluorescent/Wild-type) |  |  |  |
| --- | --- | --- | --- | --- | --- | --- |
|  |  |  | 1st eggs | 2nd eggs | 3rd eggs | Total |
| PubCas9-<br>*kmo* | 435 | 7♂ | 28/932 | 79/1334 | 31/946 | 254/8071<br>(3.147%) |
|  |  | 8♂ | 26/981 | 37/875 | 42/1268 |  |
|  |  | 12♀ | 10/620 | 7/573 | 5/542 |  |
| ExuCas9-<br>*kmo* | 350 | 7♂ | 61/869 | 61/761 | 132/1040 | 283/5934<br>(4.769%) |
|  |  | 6♂ | 0/743 | 0/311 | 0/784 |  |
|  |  | 8♀ | 2/361 | 8/540 | 6/403 |  |

Drive individuals exhibited clear red fluorescence throughout the whole body, including the larval and pupal stages, and drive homozygous individuals also showed the white eye phenotype (Fig 2a). To further verify drive insertion, we conducted PCR amplification using specific primers in the inserted construct and homology region (Fig S2a). Sanger sequencing indicted that the construct was completely inserted at the *kmo*- gRNA site (Fig S2b). To maintain these transgenic strains, we screened white-eyed mosquitoes with red fluorescence and extracted DNA from their hind legs for PCR verification. Homozygous individuals were selected to pool together, generating a long-term stable homozygous stock for subsequent experiments.

**Fig. 2.**
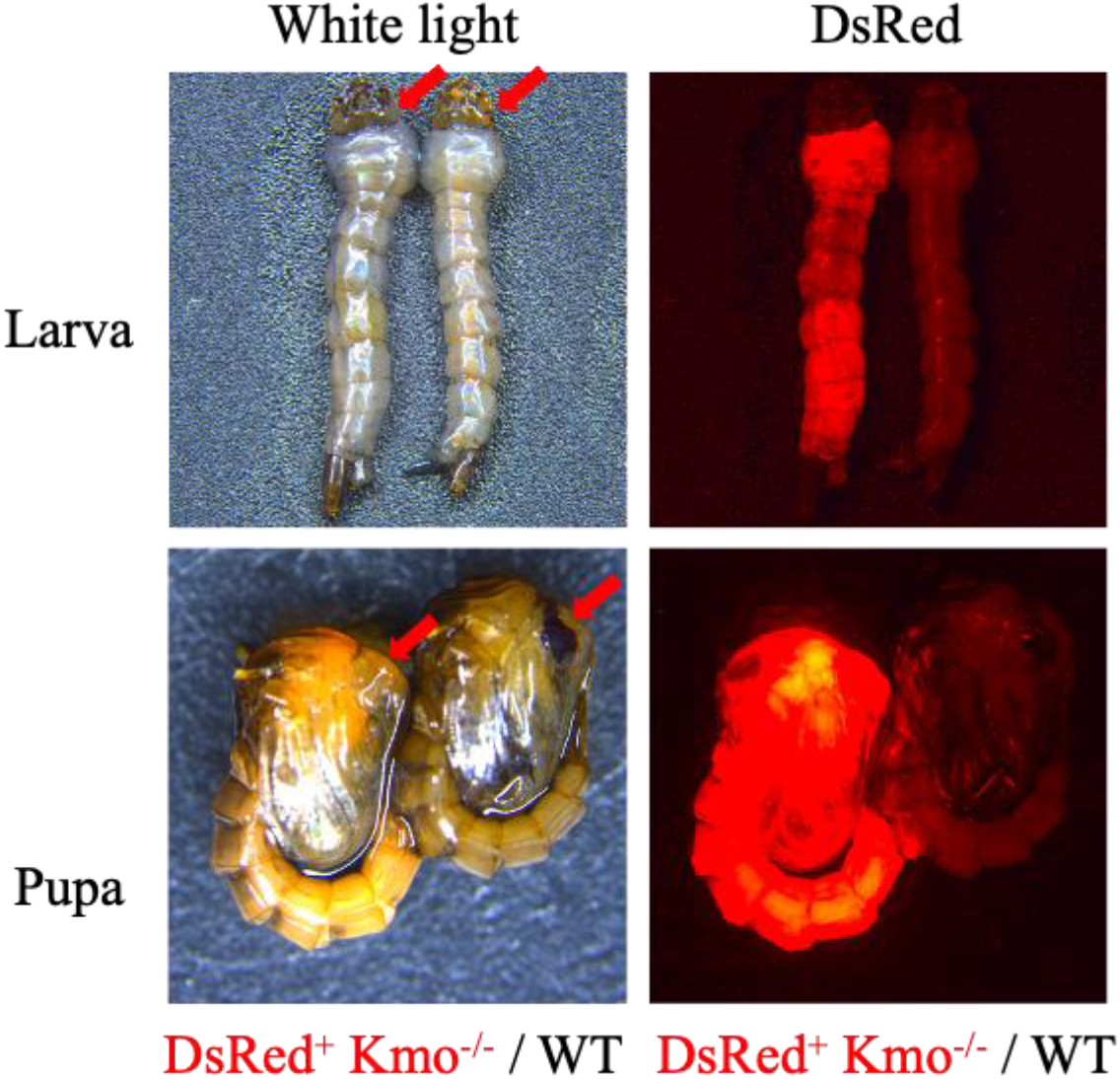
Phenotype and genotype of ExuCas9-*kmo* transgenic lines. (**a**) DsRed expression in the ExuCas9-*kmo* line. A drive homozygous mosquito is shown next to a wild-type mosquito. (**b**) PCR verification of ExuCas9-*kmo* knock-in at the *kmo* locus. Lanes 1, 4, and 7 were amplified from genomic DNA from the wild-type strain. Lanes 2, 5, and 8 were amplified from ExuCas9-*kmo* heterozygous mosquitoes. Lanes 3, 6, and 9 were amplified from ExuCas9-*kmo* homozygotes. (**c**) Sequencing map of the ExuCas9-*kmo* construct at the *kmo* locus.

### Assessment of drive efficiency

To test the drive conversion efficiency of gene drive strains, homozygous female and male mosquitoes were randomly selected to cross with wild-type. The resulting G1 mosquitoes all carry fluorescent marker genes and have two different phenotypes, DsRed fluorescent black eye (Rb) and DsRed fluorescent white eye (Rw) (Data Set S1-S2). Because there were so many white eye mosquitoes in G1 even though we did not previously detect white-eye heterozygotes, we later suspected that this effect was not due to somatic mutations. We sequenced the non-drive allele of *kmo* and found two mutations, one of which was present in all Rw drive heterozygotes. It is possible that this mutation disrupted the *kmo* gene, though it may also be linked to a mutation elsewhere that disrupts *kmo*. This mutation was ∼100 bp away from the gRNA target site, and it was linked to another mutation found ∼70 bp on the other side of the cut site, which was also found in some Rb mosquitoes (Fig S3). Each of these mutations was a single nucleotide substitution, so we did not expect that they would significantly affect drive performance. Later experiments used mosquitoes that did not have the null *kmo* mutation, but in initial testing, we separated crosses based on whether the adults were Rb or Rw.

Subsequently, male and female mosquitoes with different phenotypes of G1 were crossed to wild-type (some of which were heterozygous for the null *kmo* allele), and the G2 offspring were phenotyped (Fig 3a). Because the G1 generation were heterozygous for the drive, a Mendelian expectation of the G2 generation would be 50% red fluorescence. If the offspring exhibit a fluorescence frequency exceeding 50%, then the gene drive is likely exerting an effect.

**Fig. 3.**
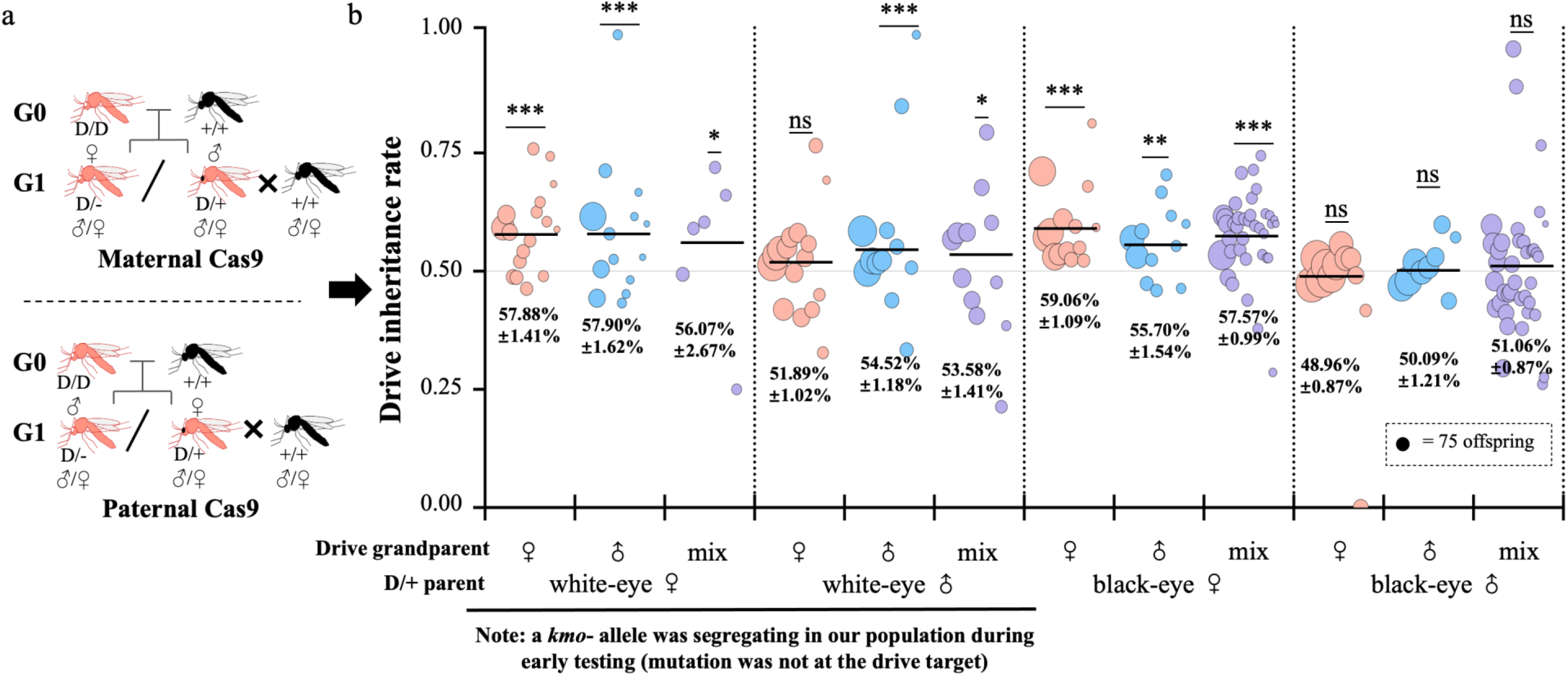
Performance of the ExuCas9-*kmo* homing gene drive. (**a**) Crossing scheme for the experiment. G0 drive homozygous grandparents were crossed to wild-type, which had some *kmo* null alleles, resulting in some white eye drive heterozygous progeny. All G1 progeny were then crossed to wild-type. (**b**) The drive inheritance rate among the G2 progeny. Data is divided to show the eye phenotype of the G1 drive heterozygous parent, and the sex of the drive homozygous grandparent is also shown. “Mix” indicates that the sex of the drive grandparent is unknown. D drive allele (with DsRed). + wild-type allele. Data shows average ± standard error of the mean. Statistical comparisons to the Mendelian expectation via the binomial test. *** *P* < 0.001, ** *P* < 0.01, * *P* < 0.05, ns – not significant.

Phenotypic screening results of the G2 generation showed that mean DsRed inheritance of PubCas9-*kmo* ranged from 43-50% for Rw and Rb females and males (Table 2 and Data Set S3). This indicated that PubCas9-*kmo* does not possess gene drive capability. All non-drive progeny has wild-type eyes, but several drive progeny had white eyes. These were not from embryo resistance or somatic activity, but were instead caused by the null *kmo* allele that was segregating in our wild-type line during this experiment.

**Table 2.**
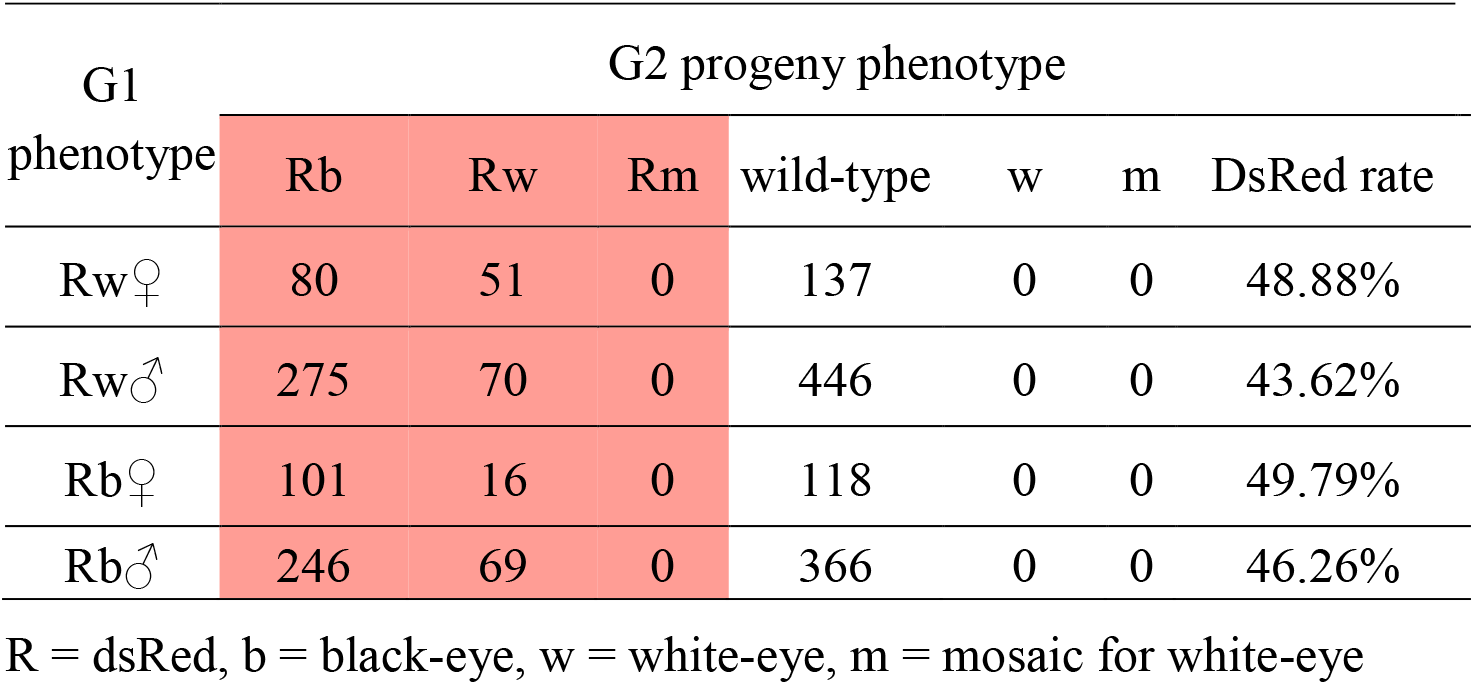
Phenotypes of progeny of G1 PubCas9-*kmo* heterozygotes crossed to wild-type.

For ExuCas9-*kmo*, we found significant differences compared to the Mendelian expectation among G2 progeny from several types of G1 crosses (Fig 3b, Table S3a and Data Set S4), though inheritance was still low compared to many other examples of homing drive and in no cases exceeded 60%. For female crosses, regardless of the eye phenotype or parent, the average transgene inheritance was 56-59%, which was significantly higher than 50% in all batches. For the male crosses, inheritance ranged from 49-55%. Only white eye males with drive fathers or with unknown drive parents had a statistically significant difference form the Mendelian expectation (Fig 3b). In no cases did we observe large differences based on drive grandparent, indicating that maternal deposition of Cas9 and gRNA did not occur at moderate to high rates. Overall, the homing efficiency of ExuCas9-*kmo* was notably higher than that of PubCas9-*kmo*. The possible activity in males of ExuCas9 was inconsistent with previous reports that shown no biased drive inheritance in male drive heterozygotes^34–37^.

To examine ExuCas9-*kmo* activity through several generations, we continued to cross different phenotypes of G2 progeny with wild-type to observe the proportion of DsRed fluorescence in G3 progeny. This was continued again to generate and phenotype G4 progeny. In these cases, flies with the same phenotype were all pooled together to set up the next set of crosses. After scoring G3 and G4 progeny, we found that transgene inheritance of drive mosquitoes with white eyes was higher than Mendelian inheritance (Table S3b, Fig S4, and Data Set S5-S6). However, black eye drive crosses observed average DsRed inheritance close to 50% without significant difference. We also noticed a gradual decrease in gene drive efficiency with successive generations. The explanation for these observations was unclear, but may be due to minor variation in batch effects or possible mosaic resistance between generations.

### Resistance allele formation in ExuCas9-*kmo*

We next sought to characterize resistance allele formation in the ExuCas9-*kmo* drive. Embryo resistance alleles form when maternally deposited Cas9 and gRNA cleaves wild-type alleles at the zygote or early embryo stage. These could be paternal alleles, but this could also happen to maternal wild-type alleles that avoid cleavage in the germline. Drive progeny with white eyes could indicate that embryo resistance allele formation took place. Somatic activity could also cause a white eye phenotype in individuals that started out as drive/Wild-type heterozygotes, via either drive conversion or resistance allele formation. Because we have identified a natural mutation in the *kmo* gene in our wild-type mosquitoes (Fig S3), accurate determination of the early embryo and somatic activity of ExuCas9-*kmo* based on phenotyping required additional observation. Assuming that the drive homozygous mosquitoes mated with black-eyed wild-type that was actually a heterozygote for the *kmo* null allele, then approximately 50% of drive offspring would have white eyes, assuming no embryo or somatic activity. We found that all crosses had close to 50% white eye progeny of those that had a drive or close to 0% white eye drive progeny (Data Set S2), indicating that only the latter were crossed to true wild-type homozygotes that that somatic/embryo activity was low. Hence, to estimate the embryo cut rate and somatic cut rate, we exclude data with over 20% white-eyed offspring. For progeny of male drive homozygotes, we found that 2.1% of drive progeny were white eye (Fig 4b and Data Set S2). This can only come from somatic activity because paternal Cas9/gRNA deposition is expected to be negligible. The final white eye rate in drive progeny of female ExuCas9-*kmo* homozygotes was approximately 1.6%, which was not significantly different from the rate in males. Thus, we can conclude that ExuCas9-*kmo* has a small amount of somatic expression, but little to no maternal Cas9 deposition.

**Fig. 4.**
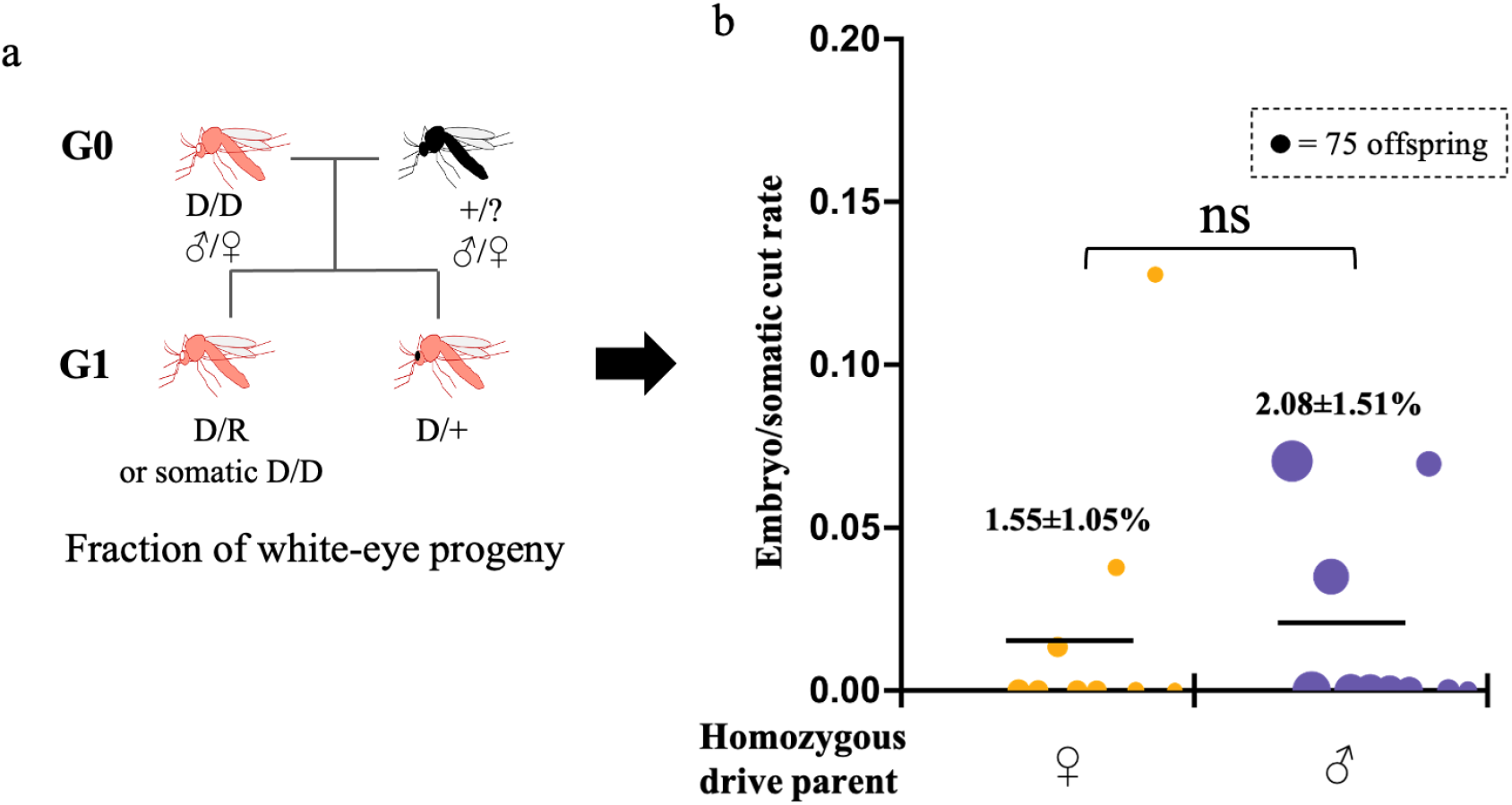
Somatic activity and resistance allele formation by maternal deposition. (**a**) Crossing scheme for investigating the embryo/somatic cut rate. Drive homozygous G0 males or females were crossed to wild-type. Crosses with ∼50% white eye G1 progeny were removed because their wild-type parent was heterozygous for the null *kmo* allele. (**b**) For the remaining crosses, the rate of white eye G1 offspring is shown as the embryo/somatic cut rate. This phenotype can come from a mix of maternally deposited Cas9 and new somatic expression of Cas9 in females, but only from somatic Cas9 expression in males. D drive allele (with DsRed). R resistance allele. + wild-type allele. Statistical comparisons between groups are via Fisher’s exact test. ns – not significant.

In order to calculate the germline resistance allele formation rate, we performed crosses between Rb mosquitoes (with male drive homozygous parents) and non-fluorescent white-eyed (w) mosquitoes that were homozygous for our *kmo* null allele (Fig 5a). We first analyzed the drive conversion rate, which was the proportion of wild-type alleles that were converted to drive alleles in the germline, assuming equal viability for all offspring genotypes. It was 13% for females and 21% for males (Fig 5b and Data Set S7). The higher than expected value for males was perhaps due to batch effects, with a few having significantly higher drive conversion than expected. To obtain the germline resistance allele formation rate, we calculated the fraction of wild-type alleles that were converted to resistance alleles. This was based on the fraction of progeny that were white eye, but lacked the drive. The germline resistance allele formation rate was 16% for drive females and 3.4% for drive males (Fig 5b and Data Set S7, a statistically significant difference (*P* < 0.001, Fisher’s exact test). Sequencing of the gRNA target site in several of these flies confirmed the presence of resistance allele sequences (Fig S5).

**Fig. 5.**
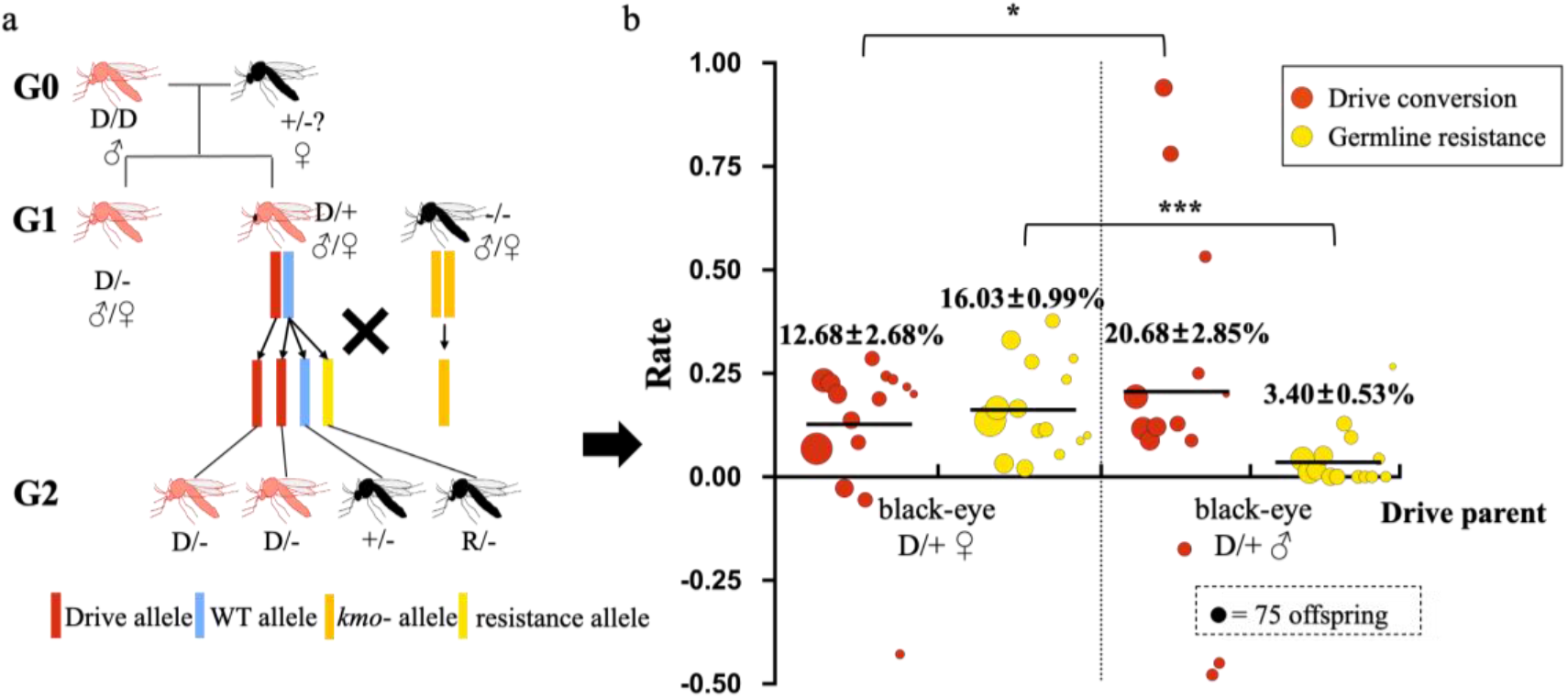
Germline resistance allele formation in the Exucas9-*kmo* drive. (**a**) Crossing scheme for analyzing germline resistance allele formation. G1 Black-eye drive heterozygotes were generated and then crossed to mosquitoes that were homozygous for the null *kmo* allele. (**b**) The germline drive conversion and resistance allele formation rates in the G1 mosquitoes based on phenotypes of G2 progeny. The germline drive conversion rate was based on the DsRed inheritance fraction. The germline resistance rate was the fraction of wild-type alleles that were converted to resistance alleles in the germline of drive heterozygotes, based on the number of non-drive progeny with white eyes. D drive allele (with DsRed). R resistance allele. + wild-type allele. - *kmo* null allele. Statistical comparisons between groups are via Fisher’s exact test. *** *P* < 0.001, * *P* < 0.05, ns – not significant.

### Improvement of microinjection efficiency using the ExuCas9-*kmo* strain

Although our lines exhibited limited gene drive effects, the ExuCas9-*kmo* line may still be suitable for gene knockout. We injected a mixture of two gRNAs and Cas9 protein into embryos, resulting in a nearly 100% mutation rate of several different target genes in G0 individuals (Table 3). Survival rates were only 3.5% to 4.5%, but mutation rates ranged from 88.89% to 100%. In comparison with direct injection into wild-type strains using mostly identical mixes (containing two gRNAs and Cas9 protein), survival of the Cas9 line was reduced, but there was a substantial increase in mutation rate achieved^47^. For the purpose of gene knockout studies, a higher mutation rate is optimal for establishing knockout strains, even at the expense of lower survival. This approach can minimize the cost and time for DNA sequencing and setting up crosses. To eliminate the introduced cas9 and gRNA, it is sufficient to selectively screen for non-fluorescent black-eyed gene knockout strains in the next generation. Therefore, the ExuCas9-*kmo* strain can be used for high efficiency gene knockout in *Ae. aegypti*.

**Table 3.** Survival and mutagenesis for the wild-type and ExuCas9-*kmo* strains.

| Strain | Target Gene | Injected G0 Embryos | G0 Survivors | G0 Mutants |
| --- | --- | --- | --- | --- |
| Wild-type | aminopeptidases<br>N 1<sup>47</sup> | 400 | 26/6.5% | 4/15% |
|  | aminopeptidases<br>N 2<sup>47</sup> | 800 | 88/11% | 36/41% |
|  | kynurenine 3-monooxygenase* | 300 | 42/14% | 3/7.1% |
|  | alkaline<br>phosphatase 1** | 500 | 78/16% | 14/18% |
|  | cadherin-like 1 | 500 | 26/5.2% | 4/5.4% |
| ExuCas9-*kmo* | aminopeptidases N 3 | 200 | 8/4% | **8/100%** |
|  | membrane-bound<br>alkaline phosphatase | 200 | 7/3.5% | **7/100%** |
|  | cadherin-like 2 | 200 | 9/4.5% | **8/88.89%** |

\*this injection mix used Cas9 mRNA instead of protein and had just one gRNA
\*\*this injection mix had just one gRNA

## Discussion

*Aedes aegypti* is a major vector mosquito and a relatively well-studied organism. The development of efficient genetic transformation technology for *Aedes aegypti* has considerably advanced research, both for basic science and for achieving environmentally-friendly control measures. Gene drive is one possible control measure, and here, we characterized a complete gene drive using the well-studied *exu* promoter for Cas9. Drive inheritance was low, but usually still super-Mendelian. As expected, the embryo resistance allele formation rate was low (if germline expression was low, there should be little Cas9 available to maternally deposit). We observed some somatic expression, and germline resistance rates were similar to drive conversion rates.

In recent years, the CRISPR/Cas9 nuclease system has been widely used in *Aedes aegypti* functional genetics research^46, 49^. However, there is still room for improvement in terms of enhancing the stability and efficiency CRISPR/Cas9 editing, particularly with regards to facilitating genetic knock-in. It is possible to effectively enhance the homology-directed repair mechanism for knock-in by inhibiting *lig4* and *ku70*, which mediate the non-homologous end-joining mechanism^50, 51^. In human and mouse cell lines, the efficiency of CRISPR/Cas9-mediated HDR was increased eightfold using SCR7 inhibitor or reducing the expression levels of *lig4* and *ku70* through RNAi^51^. The addition of *ku70*-inhibiting dsRNA in Cas9-sgRNA injection also raised the gene knock-in rate of *Drosophila melanogaster*^52^, *Bombyx mori*^50^ and *Aedes aegypti*^53^. In order to improve the likelihood of obtaining transgenic *Aedes aegypti*, we also added ds-*ku70* to the injection solution, mixing Cas9 protein, *kmo*-gRNA, Cas9-*kmo* plasmid, and *ku70*-dsRNA. Consequently, our gene knock-in rates reached 3.1% and 4.8%. However, whether the dsRNA affected the efficiency of *Ae. aegypti* knock-in still needs systematic verification.

Due to the inclusion of multiple expression elements, gene drive systems can be over 10,000 bp, making it extremely challenging to achieve targeted genome integration. The success rates of CRISPR/Cas9 knock-in in the G1 generation ranged from 0.05% to 0.38% in *An. Stephensi*, while only two male individuals expressing endogenous CRISPR/Cas9 were found among 25,712 G1 larvae in *An. Stephensi*^10^. Another recent study got 96 successful transformants from 25,296 G1 larvae in *An. Stephensi*^32^. Similarly, only 21 transgenics were obtained among 5906 G1 larvae with 0.36% success rate for ∼15.6 kb drive element insertion in *An. gambiae*^31^. Therefore, CRISPR/Cas9-mediated long fragment gene knock-in demonstrates significantly higher genetic transformation efficiency in *Ae. aegypti* compared to *Anopheles* mosquitoes. This is somewhat incongruous with the substantially higher drive efficiency observed in *Anopheles*^10, 11, 30–32^ compared to *Aedes*^35–37^. However, endogenously expressing *Aedes* Cas9 trains showed good potential as a tool for gene knockout^42^. When gRNA only was injected into Cas9 strain eggs, the gene editing rate in G0 survivals ranged from 29% to 94%, with Exu-Cas9 having the best editing effect of 61-87%^42^. When we injected gRNA into eggs of our Exu-Cas9 strain, additional Cas9 protein was added, which combined with the genomic insertion site and possibly injection technique allowed us to consistently improve our G0 mutagenesis rate with three different targets to 100%.

Currently, the remarkable progress of CRISPR/Cas9 has led to extensive investigations into gene drive systems for genetic regulation of pest populations. Similar to gene drive systems in fruit flies^54^ *An. gambiae*^11^, and *An. Stephensi*^10^, the gene drive system developed in this study for *Ae. aegypti* uses a complete drive rather than split drive. The promoter of Cas9 expression is one of the key elements in gene drive systems, determining the efficiency of Cas9 cleavage in germ cells. In this study, we tested two endogenous promoters from *Ae. aegypti*, the ovary-specific high-expression promoter *Exu* and the ubiquitous promoter *Pub*^42^. The drive allele inheritance rate of ExuCas9- *kmo* was low but super-Mendelian, while pubCas9-*kmo* had no driving effect. So far, there are four other studies describing the development of gene drives in *Ae. aegypti*, evaluating the driving efficiency of various promoters^34–37^. Most of these gene drives are split drives, and the strains expressing Cas9 were generated using the PiggyBac transposase system^38, 55^. Owing to the uncertainty of PiggyBac insertion sites, the same promoter-Cas9 combination may result in different lines with varying insertion sites and different Cas9 expression. This can produce completely different genetic environments, leading to varied Cas9 expression and gene drive efficiency. For example, for the same *sds3* promoter, inheritance rates could reach over 80%, while other lines were close to 50%^35^. Similarly, for the *shu* promoter, inheritance rates varied from 50% to 90% among four different strains^35^. Therefore, split drives may not be completely reflective of the possibilities for a complete drive. Though some combinations gave higher efficiency, the only other study that made complete drives had relative low drive inheritance with the *nanos* and *zpg* promoters^38, 55^.

Similarly, the choice of gRNA promoter significantly influences gene drive efficiency. We used a single 400 bp U6-1 promoter sequence, together with a 300 bp terminator. At the same time of this study, Li et al. employed six 1000 bp U6 promoters from *Ae. aegypti* to construct gRNA expression vectors^34^. These vectors were injected into embryos of exuCas9 and ub40Cas9 strains, causing gene mutations in the injected mosquitoes. Among them, U6b (AEEL017774, our U6-1) and U6c (AEEL017763) exhibited higher activity. It is possible that our 400bp AeU6-1 promoter had lower activity than the 1000 bp version, account for our somewhat lower drive activity^34^, though this needs further verification.

Our discovery of a possible single nucleotide C:T *kmo* null mutation was most likely a natural mutation. If confirmed to be the cause of the white eye phenotype, this could potentially provide a promising target site for development single-base editing techniques in *Ae. aegypti*.

In summary, our exuCas9-kmo strain in *Ae. aegypti* provides useful data on gene drive and gene editing efficiency. The utilization of this strain may greatly enhance the success rate of gene editing experiments in *Ae. aegypti* and provide robust technical support for future functional genetics research. The analysis of this standard gene drive system in *Aedes aegypti* showed low drive conversion efficiency and relatively high resistance allele formation, both of which will need to be overcome in this species to further expand the potential of gene drive technology to develop efficiency and environmentally friend pest control tools..

## Acknowledgements

This project supported by the Science Foundation of Fujian Province, China grant number 2023J011576 to JW, National Science Foundation of China grant number 32302455 to XX, and National Science Foundation of China grant number 32270672 to JC, who also received support from the Center for Life Sciences.

## Supplemental Information

**Table S1.**
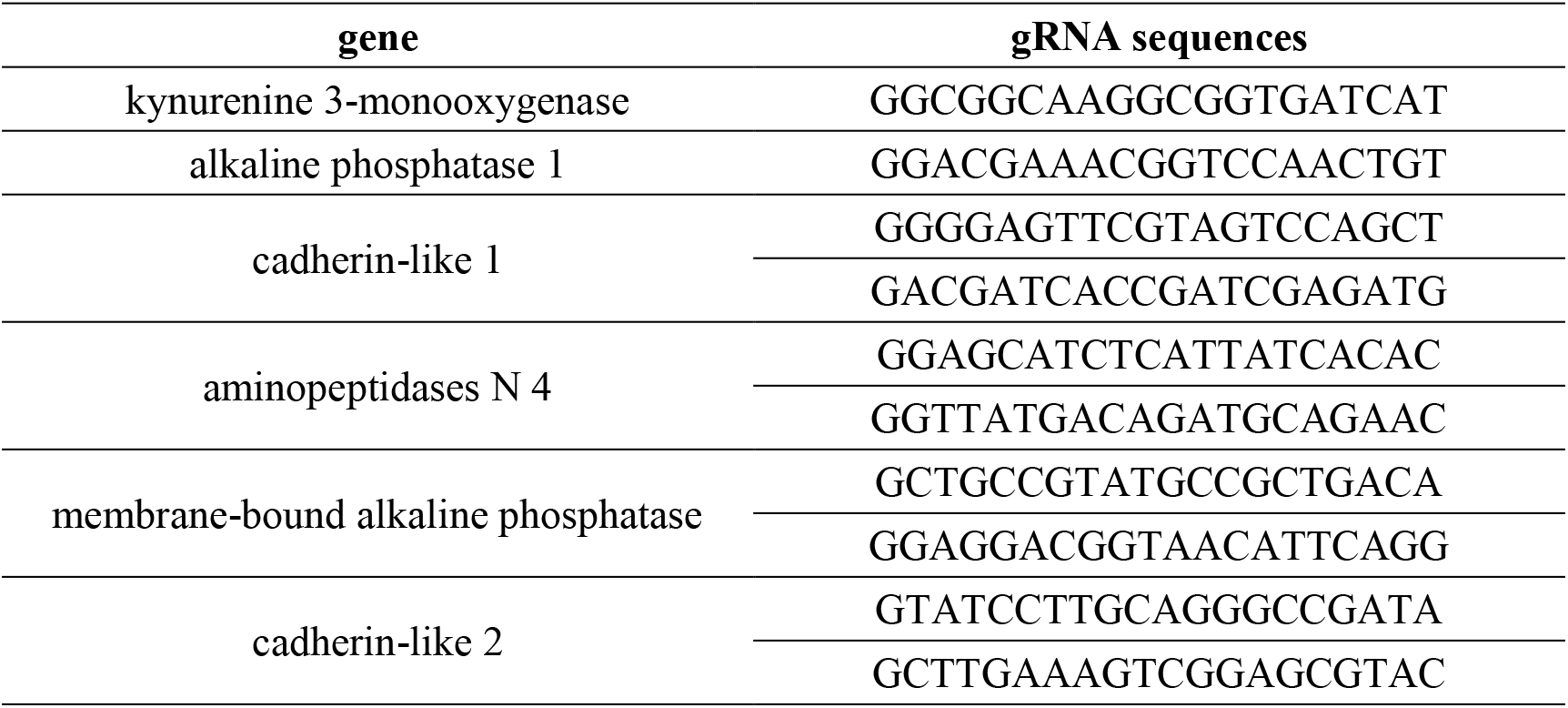
Sequences of gRNA.

**Table S2.**
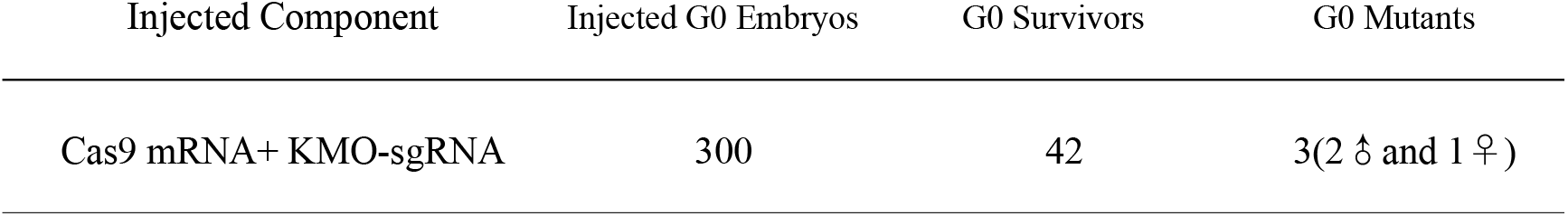
Summary of the survival and mutagenesis rates from injected *kmo*-sgRNA.

| Injected Component | Injected G0 Embryos | G0 Survivors | G0 Mutants |
| --- | --- | --- | --- |
| Cas9 mRNA+ KMO-sgRNA | 300 | 42 | 3(2 ♂ and 1 ♀) |

**Fig. S1.**
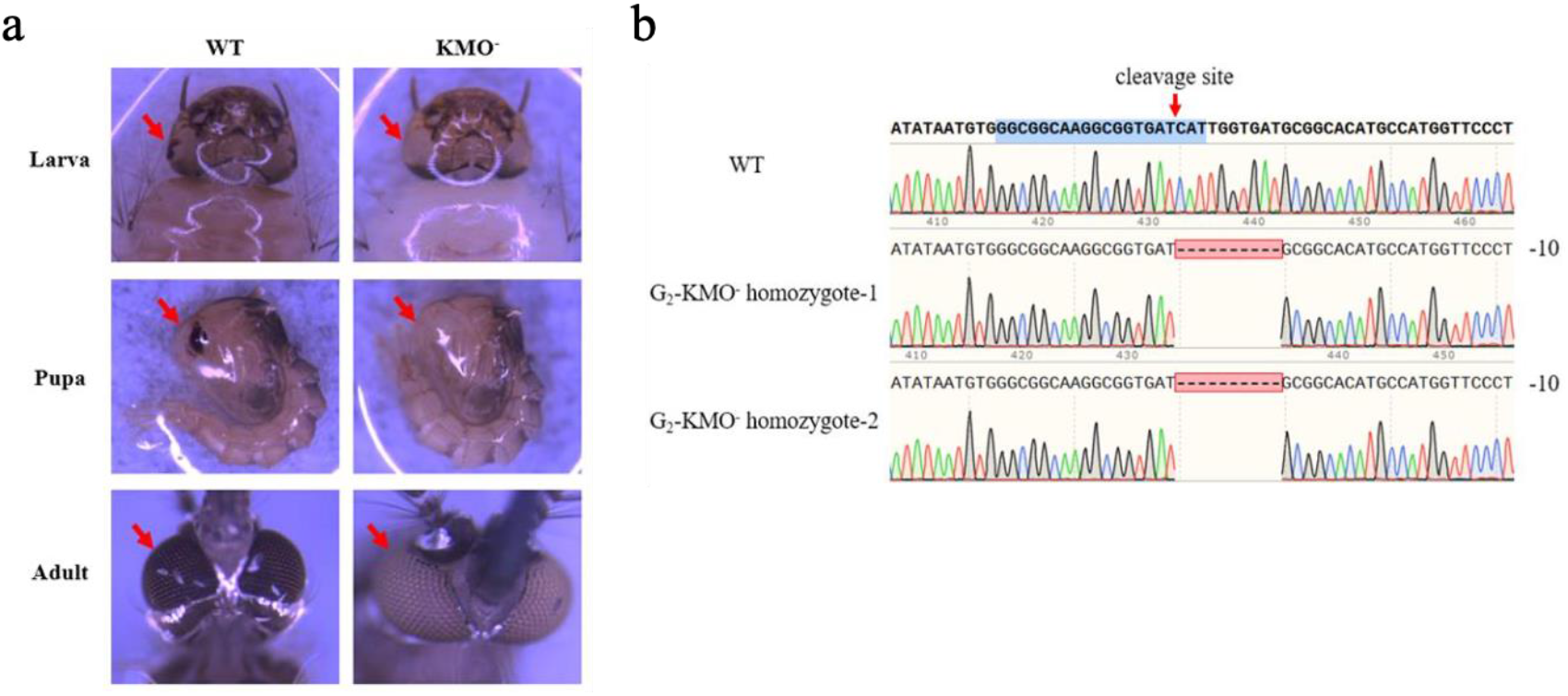
*Ae. aegypti kmo*^-^ strain. (**A**) CRISPR/Cas9-generated white-eye phenotype of mutant *Ae. aegypti* strains. (**B**) DNA sequencing chromatograms of the CRISPR/Cas9 target site region from a G_0_ white-eye mosquito. The red arrow shows the Cas9 cleavage site.

**Fig. S2.**
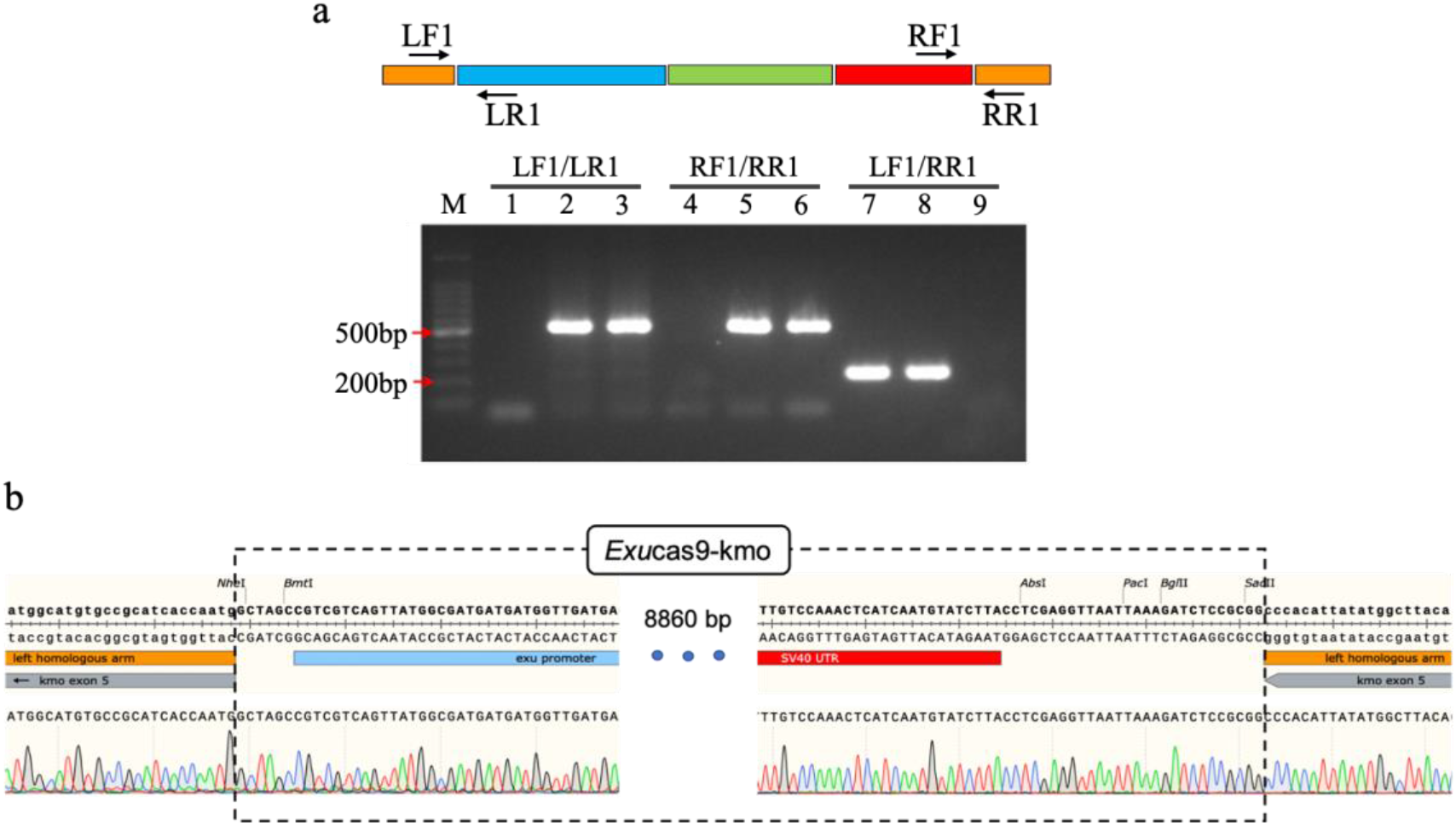
Genotype of ExuCas9-*kmo* transgenic lines. (**a**) PCR verification of ExuCas9-*kmo* knock-in at the *kmo* locus. Lanes 1, 4, and 7 were amplified from genomic DNA from the wild-type strain. Lanes 2, 5, and 8 were amplified from ExuCas9-*kmo* heterozygous mosquitoes. Lanes 3, 6, and 9 were amplified from ExuCas9-*kmo* homozygotes. (**b**) Sequencing map of the ExuCas9-*kmo* construct at the *kmo* locus.

**Fig. S3.**
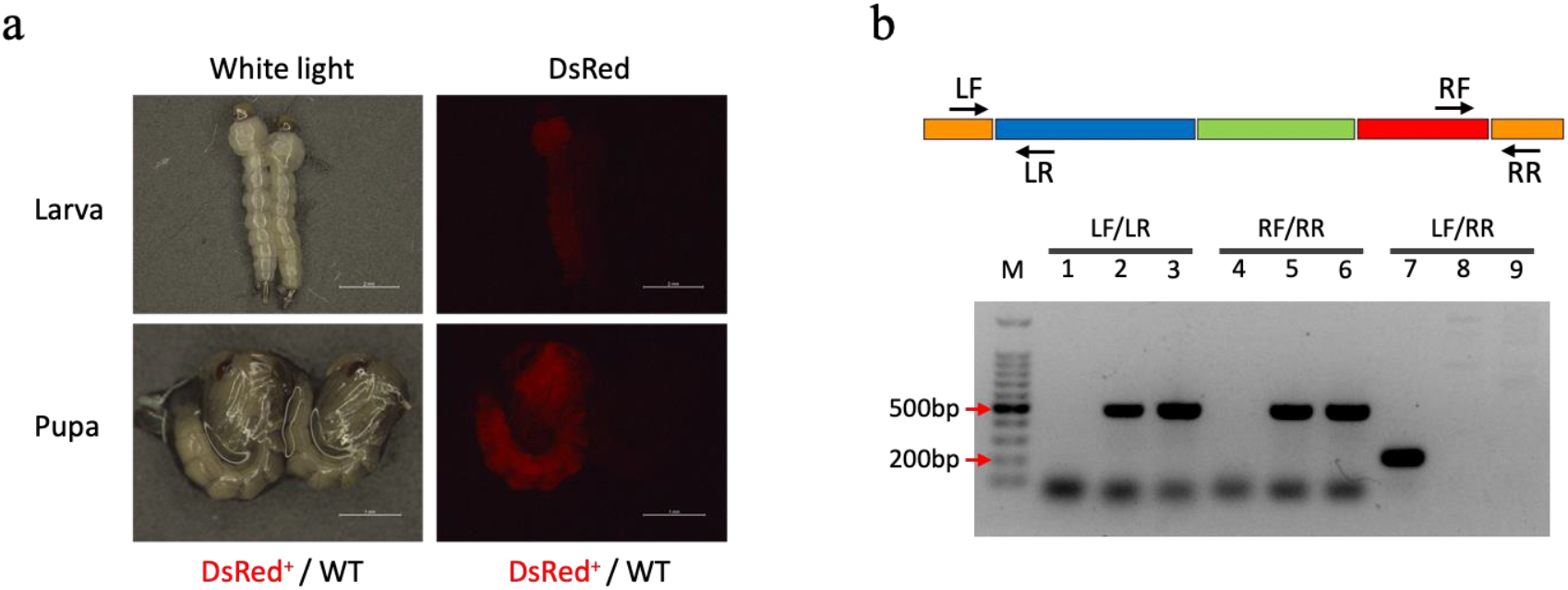
Phenotype and genotype of PubCas9-*kmo* transgenic line. (**a**) Microscopy under white light and imaging DsRed for a heterozygous PubCas9-*kmo* larva and a wild-type larva. (**b**) PCR verification of the PubCas9-*kmo* knock-in. Lanes 1, 4 and 7 were amplified from genomic DNA of the wild-type strain. Lanes 2, 5 and 8 were amplified from female PubCas9-*kmo* homozygotes. Lanes 3, 6 and 9 were amplified from male PubCas9-*kmo* homozygotes.

**Fig. S4.**
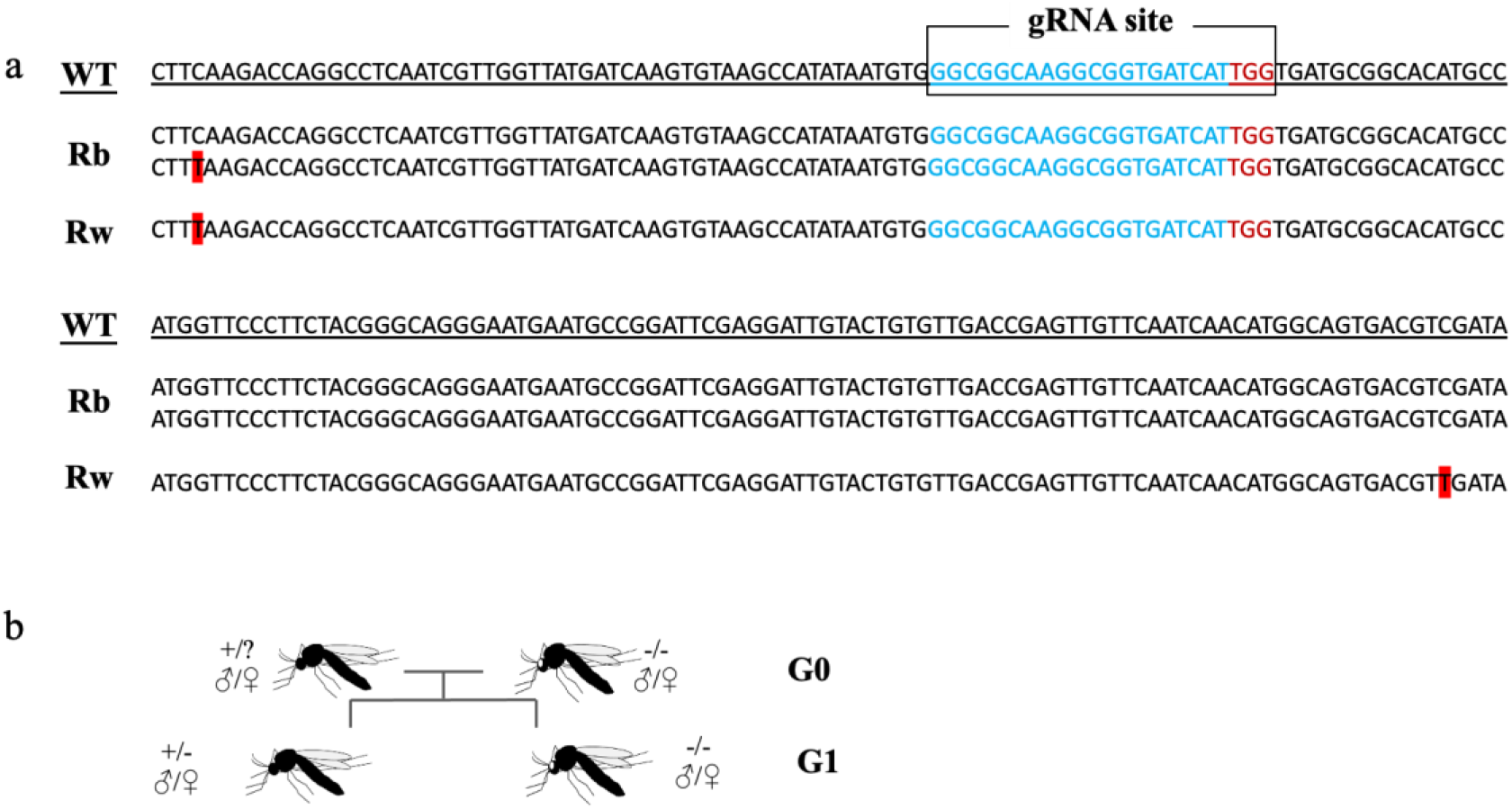
Sequences of the *kmo* gene near the drive target site. (**a**) DNA sequence of the wild-type *kmo* gene around gRNA target site, together with specific sequences from the non-drive allele in drive flies with black (wild-type) or white eyes. Red highlighted nucleotides are mutations from wild-type. Note that two separate sequences were obtained from Rb flies, one of which was identical to the expected reference sequence. (**b**) Crossing scheme for obtaining mutations in the *kmo* gene. White-eye progeny from the cross had a mutation.

**Table S3.** Assessment of ExuCas9-*kmo* drive.

a: Phenotypes of the progeny of G1 mosquitoes
| G1 phenotype | R | Rw | Rm | wild-type | w | m | DsRed rate |
| --- | --- | --- | --- | --- | --- | --- | --- |
| Rw♀ | 1073 | 361 | 0 | 974 | 80 | 0 | 57.64% |
| Rw♂ | 2362 | 540 | 0 | 2382 | 177 | 0 | 53.14% |
| Rb♀ | 1920 | 1253 | 58 | 2173 | 185 | 4 | 57.77% |
| Rb♂ | 2956 | 1198 | 0 | 4121 | 29 | 0 | 50.02% |

b: Phenotypes of the progeny of G2 mosquitoes
| G2 phenotype | R | Rw | Rm | wild-type | w | m | DsRed rate |
| --- | --- | --- | --- | --- | --- | --- | --- |
| Rw♀ | 705 | 124 | 0 | 656 | 24 | 0 | 54.94% |
| Rw♂ | 1689 | 238 | 0 | 1436 | 5 | 0 | 57.21% |
| Rb♀ | 727 | 99 | 0 | 868 | 1 | 0 | 48.73% |
| Rb♂ | 1306 | 285 | 0 | 1492 | 2 | 0 | 51.57% |

c: Phenotypes of the progeny of G3 mosquitoes
| G3 phenotype | R | Rw | Rm | wild-type | w | m | DsRed rate |
| --- | --- | --- | --- | --- | --- | --- | --- |
| Rw♀ | 942 | 72 | 0 | 872 | 0 | 0 | 53.76% |
| Rw♂ | 1296 | 92 | 0 | 1236 | 0 | 0 | 52.90% |
| Rb♀ | 815 | 236 | 0 | 995 | 12 | 0 | 51.07% |
| Rb♂ | 1229 | 81 | 0 | 1375 | 0 | 0 | 48.79% |

R = dsRed, b = black-eye, w = white-eye, m = mosaic for white-eye

**Fig. S5.**
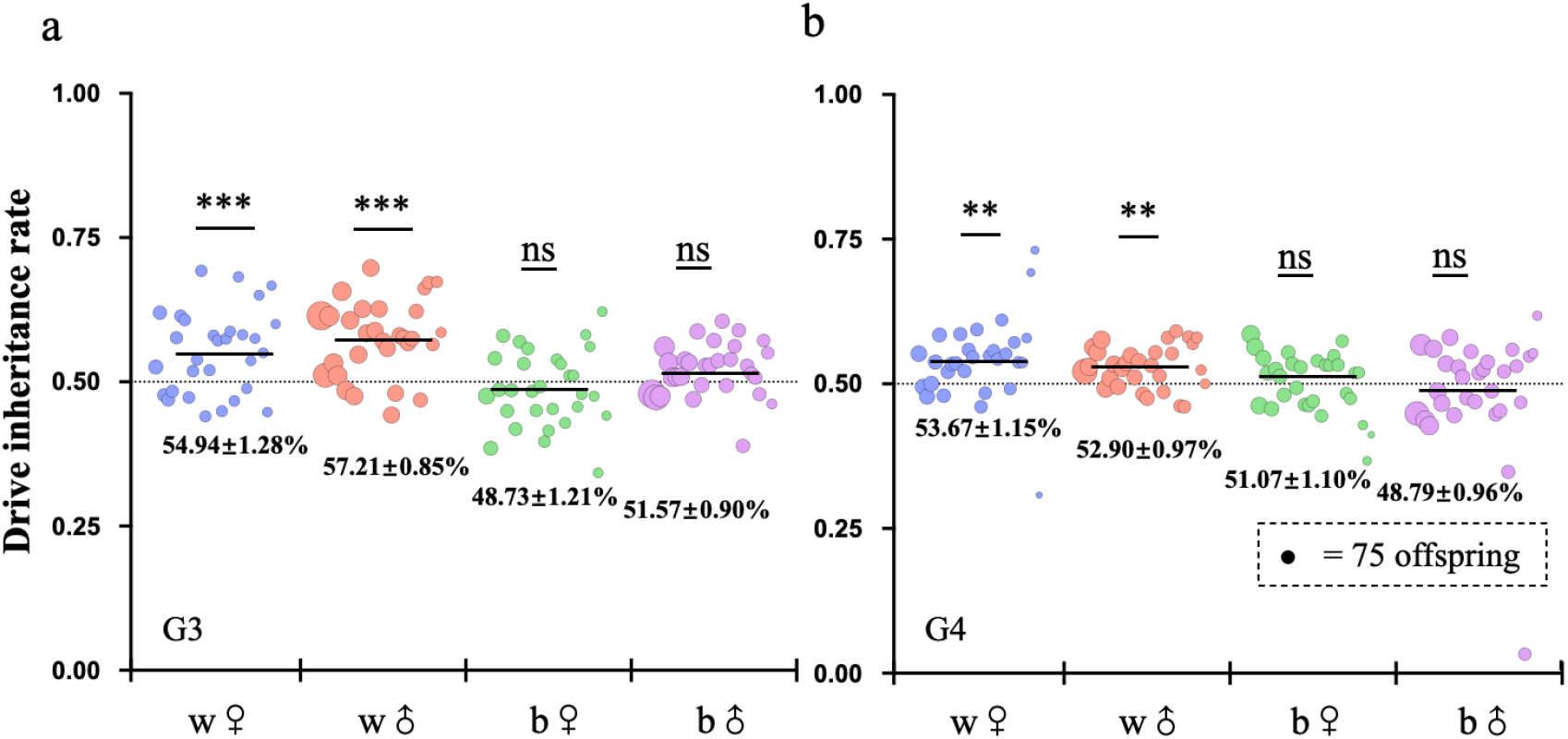
Drive inheritance in later generations from Exucas9-*kmo* homing drive crosses. The chart shows the inheritance rates in the (**a**) G3 and (**b**) G4 offspring generated from crosses between wild-type and G2 or G3 mosquitos, respectively. The horizontal position shows the eye phenotype of the drive parent. w: white-eye. b: black-eye (wild-type). Data shows average ± standard error of the mean. Statistical comparisons to the Mendelian expectation via the binomial test. *** *P* < 0.001, ** *P* < 0.01, ns – not significant.

**Fig S6.**
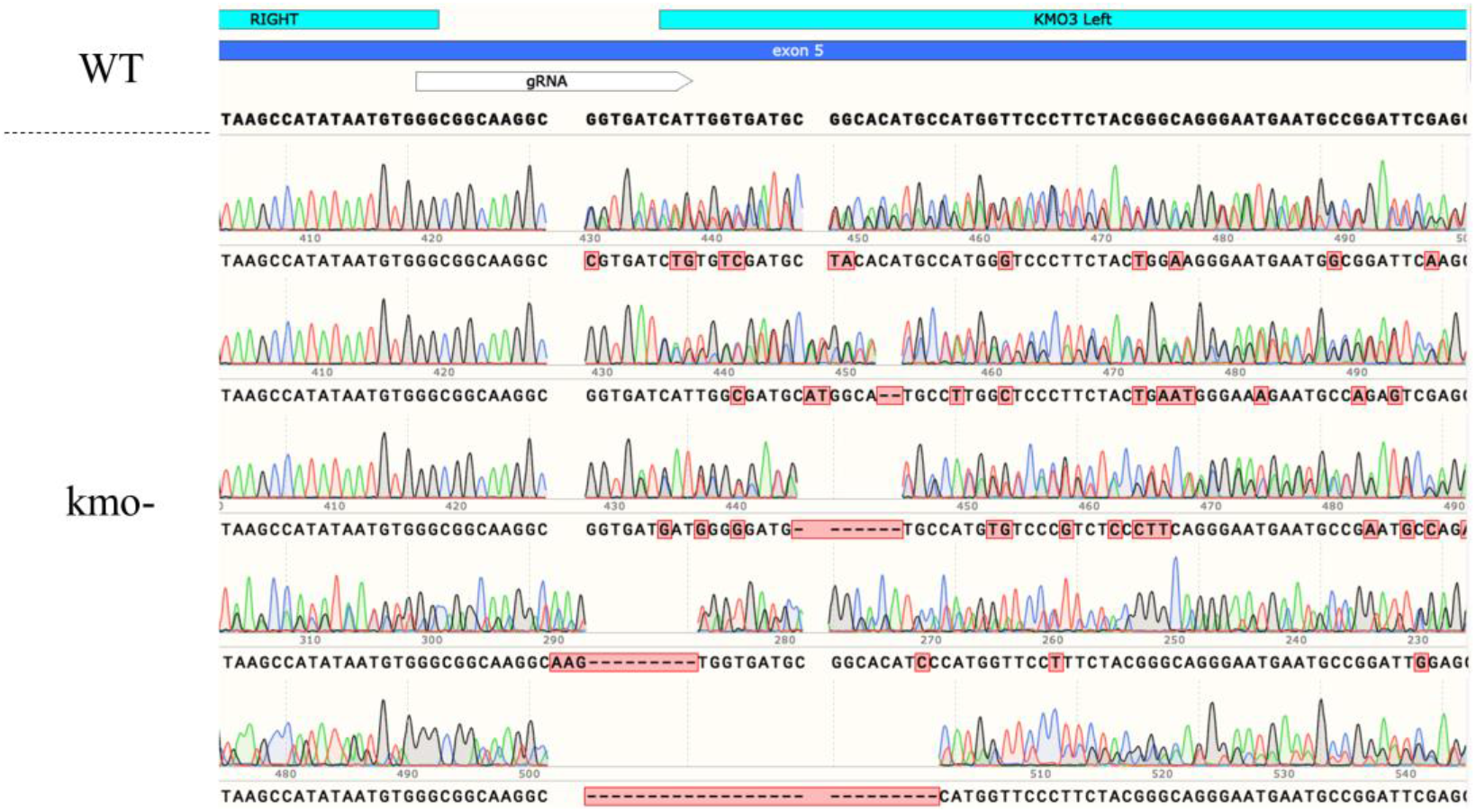
Sequencing of resistance alleles. Sequences from non-drive progeny of drive individuals that had resistance alleles. Sequences are mosaic because they also contain a wild-type allele from the other parent.

## Notes

https://github.com/jchamper/Aedes-Complete-Drive

### Competing Interest Statement

The authors have declared no competing interest.

